# Transcriptomic and rRNA:rDNA signatures of environmental vs. enteric *Enterococcus faecalis* isolates under oligotrophic freshwater conditions

**DOI:** 10.1101/2021.05.04.442698

**Authors:** Brittany Suttner, Minjae Kim, Eric R. Johnston, Luis H. Orellana, Carlos A. Ruiz-Perez, Luis M. Rodriguez-R, Janet K. Hatt, Joe Brown, Jorge W. Santo Domingo, Konstantinos T. Konstantinidis

## Abstract

The use of enterococci as a fecal indicator bacterial group for public health risk assessment has been brought into question by recent studies showing that “naturalized” populations of *E. faecalis* exist in the extraenteric environment in a viable but not culturable (VBNC) state. The extent to which these naturalized or VBNC *E. faecalis* can confound water quality monitoring is unclear. To determine if strains isolated from different habitats display different survival strategies and responses, we compared the decay patterns of three *E. faecalis* isolates from the natural environment (*environmental strains*) against three human gut isolates (*enteric strains*) in laboratory mesocosms that simulate an oligotrophic, aerobic freshwater environment. Our results showed similar overall decay rates between enteric and environmental isolates based on viable plate and qPCR counts. However, the enteric isolates exhibited a spike in rRNA:rDNA ratios between days 1 and 3 of the mesocosm incubations that was not observed in environmental isolates, which could indicate a different stress response. Nevertheless, there was no strong evidence of differential expression of genes thought to be related to habitat adaptation in the accompanying mesocosm metatranscriptomes when compared between environmental and enteric isolates. Overall, our results provide novel information on how rRNA levels may vary over different metabolic states (i.e., alive vs. VBNC) for this important indicator bacteria. We also observed some evidence for habitat adaptation in *E. faecalis*; however, this adaptation may not be substantial or consistent enough for integration in water quality monitoring.

**IMPORTANCE:** Enterococci are commonly used worldwide to monitor environmental fecal contamination and public health risk for waterborne diseases. However, some species within this group can enter an inactive, viable but not culturable (VBNC) state that make it difficult to accurately quantify during routine monitoring. Furthermore, lower-risk, environmental enterococci strains may also confound water quality estimates. We developed an rRNA:rDNA viability assay for *E. faecalis* (a predominant species within this fecal group) and tested it against both enteric and environmental isolates in freshwater mesocosms to assess whether this approach can serve as a more sensitive water quality monitoring tool. We were unable to reliably distinguish the different isolate types using this assay under the conditions tested here; thus, environmental strains should continue to be counted during routine water monitoring. However, this assay could be useful for distinguishing more recent (i.e., higher risk) fecal pollution because rRNA levels significantly decreased after one week in all isolates.

## INTRODUCTION

Enterococci are used worldwide as fecal indicator bacteria based on the assumptions that they are predominantly found in intestinal systems of animal hosts and exhibit high die-off rates upon release to the natural environment. However, populations of *Enterococcus* spp. have been isolated from freshwater environments with no sign of recent fecal inputs (heretofore referred to as “environmental” strains) (1, 2). These environmental strains are phenotypically and phylogenetically indistinguishable from their enteric relatives based on standard selective media so their recovery during a water quality test by conventional methods would be considered a positive indicator of fecal contamination (3, 4). Whole genome comparisons of environmental and enteric strains have revealed distinct habitat-specific genetic signatures. For example, enteric genomes were specific or highly enriched for genes associated with antibiotic resistance, virulence, and the metabolism of sugars, while nickel and cobalt transport systems were overrepresented in the environmental genomes (4–6). These results suggested that the accessory gene content carried by different *E. faecalis* isolates may contribute to differential survival and adaptation in different habitats despite high genetic relatedness among core genes present in all isolates (4, 6). However, the practical application and use of these alternative gene markers to distinguish innocuous environmental strains from enteric strains that indicate a risk to public health have not yet been tested.

*Enterococcus faecalis* is one of the most abundant and often isolated bacterial species within the fecal enterococci group. Moreover, species-specific qPCR studies have shown their prevalence in different animal fecal sources and in environmental waters (7). *E. faecalis* is known to enter a viable but non-culturable (VBNC) state as a survival response to environmental stressors, such as introduction into an extraenteric environment (8). VBNC cells preserve membrane integrity and low levels of gene expression, but typically do not form colonies using traditional culture-based methods and have distinct proteomic signatures compared to non-VBNC cells (9, 10). However, VBNC cells can be resuscitated and grow upon return to favorable conditions (11), and thus for pathogenic bacteria, VBNC cells may represent risks to public health while VBNC fecal indicator bacteria may be relevant to exposure risks. Accordingly, culture-based approaches can lead to inaccurate assessments of health risks due to VBNC (false negatives) or natural reservoirs of enterococci (false positives). Elucidating the extent to which naturalized populations and/or VBNC cells may confound water quality monitoring is therefore critical for robust public health risk assessment.

Several studies have used cellular ribosomal RNA levels, often expressed as the copy number ratio of 16S rRNA gene transcripts to 16S rRNA gene DNA copies (i.e., rRNA:rDNA ratio), to detect active and/or growing microbes (12–15). This is based on the assumption that the levels of ribosomal RNA are much higher in actively growing and metabolizing cells relative to VBNC (8), dormant, or dying cells. Further, examining the rRNA:rDNA ratio level may be a more accurate assessment of the state of cellular activity than techniques based on membrane permeability (e.g., PMA-qPCR or live-dead staining microscopy) because cell death can occur before cell membrane lysis (16). Although the usefulness of the rRNA:rDNA ratio for these purposes has been documented for several bacterial genera, the relationship between rRNA:rDNA ratio and growth rate varies significantly between taxa and some studies have even reported an inverse relationship between rRNA concentrations and growth rate (17–19). Since these ratios appear to be, at least partly, taxon-specific, baseline data on rRNA:rDNA levels in *E. faecalis* during different stages of activity and decay are needed in order to determine if it can be used as a viability assay for water quality monitoring to distinguish environmentally adapted from enteric strains.

Accordingly, the guiding hypothesis of this study is that the strains associated with different habitats (i.e., enteric vs. environmental) have distinct genetic and/or physiological adaptations that cause differential survival in freshwater ecosystems, and this can be detected and quantified for more accurate public health risk assessment based on rRNA:rDNA gene copy number ratios and whole-genome gene expression profiles. In particular, it is expected that environmentally-adapted *E. faecalis* strains (if such strains exist) would have higher rRNA:rDNA ratios in surface water environments compared to enteric strains because the former strains are better able to survive environmental stressors like O_2_, sunlight, and nutrient limitation (20). In contrast, enteric strains, if they can persist in that same environment, are expected to be in a lower activity state.

To test this hypothesis, we performed laboratory mesocosm incubations that simulated the natural freshwater environment with three environmental and three enteric *E. faecalis* isolates that were previously reported to be phylogenetically and phenotypically indistinguishable from one another (4). The change in viable cell counts (i.e., plate counts) and rRNA:rDNA ratios were then monitored over two weeks to assess their decay and metabolic state. Therefore, this study provides important baseline information on rRNA levels in *E. faecalis* under different growth conditions and new insights into the use of rRNA for improved water quality monitoring.

## RESULTS

### *E. faecalis* strains used in the mesocosm incubations

Individual *E. faecalis* isolates (3 enteric and 3 environmental; Table 1) were selected based on previous comparative genomic analysis that showed these isolates contain putative habitat-specific gene signatures (4). The environmental isolates were recovered from freshwaters with unknown history of fecal pollution sources using standard selective medium for enumerating enterococci. The enteric isolates were recovered from the human GI tract and were publicly available as part of the Human Microbiome Project (21). All of these six strains shared >97% average nucleotide identity (ANI), well above the 95% ANI cutoff used for species demarcation (22). Further details on the source and genome content of these isolates are described in Weigand et al. (4).

**Table 1:** *E. faecalis* isolates used in the dialysis bag mesocosm experiments. Total RNA from the mesocosm samples was also analyzed with metatranscriptomics for the isolates in bold.

| **Isolate Name** | **Isolation source<sup>a</sup>** | **GenBank Accession** |
| --- | --- | --- |
| MMH594 | enteric | AJDZ01000001.1 |
| **ERV62** | enteric | ALZQ01000001.1 |
| **TX0104** | enteric | ACGL01000001.1 |
| MTUP9 | environmental | AYOJ01000001.1 |
| **MTmid8** | environmental | AYKU01000001.1 |
| **AZ19** | environmental | AYLU01000001.1 |

<sup>a</sup>Isolation source describes whether the strains were isolated from the human gut (enteric) or freshwaters with unknown history of fecal pollution sources (environmental) according to Weigand et al., 2014.

### rRNA:rDNA ratio of enteric versus environmental *E. faecalis* isolates in dialysis bag mesocosms simulating an oligotrophic freshwater habitat

Oligotrophic growth conditions invariably induced the VBNC state for both enteric and environmental strains of *E. faecalis* in our pilot experiment using glass bottle mesocosms based on viable cell vs. qPCR counts (see supplemental material for details). However, in this experiment using only traditional qPCR of the 16S rRNA gene, we were not able to measure any differences in cellular viability (i.e., live, dead, or VBNC) between the two strain types (Figure S1). Therefore, we subsequently used the rRNA:rDNA ratio to compare the physiological response of human enteric versus environmental *E. faecalis* isolates in laboratory dialysis bag mesocosms simulating an oligotrophic freshwater environment. Specifically, *E. faecalis* isolates were spiked in known concentrations in filter-sterilized lake water that was enclosed in dialysis bags, and the bags were subsequently incubated in 10-gallon aquarium tanks filled with (unfiltered) lake water. Dialysis bags allow water and nutrients to pass through but not cells and thus, represent convenient systems for incubations that simulate well *in situ* conditions. Mesocosm sampling occurred on days 1, 3, 8 and 11 (D1, D3, D8 and D11) and included plating for viable cell counts and filtering for total nucleic acid extraction to determine rRNA:rDNA ratios. Although the lake water used in the dialysis bags was filter-sterilized before inoculation (i.e., to remove predators), some growth was observed in water from the negative control bags by D8 on tryptic soy agar (TSA) plates. However, no growth was observed on the *Enterococcus-*specific media (data not shown). This result suggested that the integrity of the dialysis bags started to break down over time and that some of the microbes from the non-sterilized lake water in the tanks outside of the bags were able to pass through the dialysis membranes. Hence, we primarily focused our analysis and interrelations on the first three sampling points.

All strains exhibited a decrease in viable cell counts for the duration of the experiment (i.e., until D11; Figure 1A). Moreover, decay rates based on plate counts were not significantly different between the two isolate types, i.e., *environmental* vs. *enterics* (paired Wilcoxon P=0.063), consistent with our previous pilot experiment (Figure S1). The average rRNA:rDNA ratios in the three environmental and one of the enteric isolates were relatively stable from D1 to D3 (0.7 to 1.5-fold change in the ratio). However, two of the enteric strains (ERV62 and MMH594) had an ~6-fold increase in their rRNA:rDNA ratios from D1 to D3 (Figure 1B). By D8 and D11, the average ratios decreased and approached zero consistently for all six isolates. When comparing average rRNA:rDNA ratios overall (i.e., enteric vs. environmental across all time points), the enteric isolates were not significantly different from environmental isolates (Wilcoxon Rank Sum P=0.149). When looking at the habitat types separately (i.e., among themselves) over time, the average rRNA:rDNA ratios between D1 and D3 were not significantly different for the three environmental strains but were significant for the three enteric strains (paired Wilcoxon P= 1.0 and 0.014, respectively). This result suggested that the enteric and environmental isolates may show different gene expression responses to environmental stress (e.g., nutrient limitation), which we examined more fully with metatranscriptomics.

**Figure 1:**
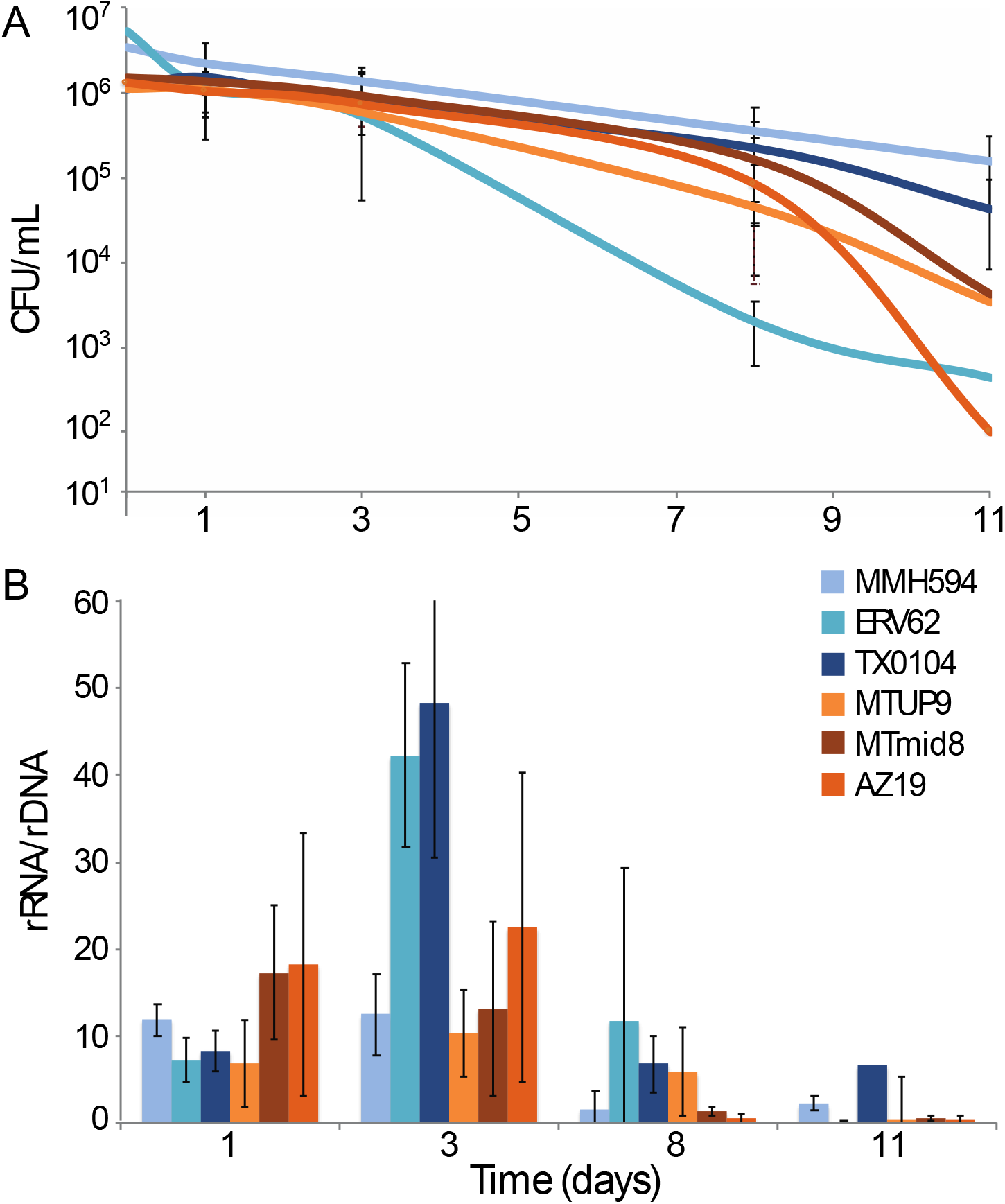
Comparing changes in **(A)** viable cell counts and **(B)** rRNA:rDNA ratios of enteric versus environmental *E. faecalis* isolates over time in dialysis bag mesocosms. Three enteric and three environmental isolates are represented by different shades of orange and blue, respectively. Error bars are standard deviation among three technical replicates.

### Comparative metatranscriptomics of enteric and environmental isolates

The 16S rRNA:rDNA ratio alone does not provide information about specific gene functions and differences in mRNA expression levels that may serve as more sensitive biomarkers for the response to oligotrophic freshwater conditions. Thus, we used metatranscriptome sequencing profiles of the dialysis bag mesocosms to identify specific metabolic pathways that may underlie habitat adaptation and represent more reliable targets for improved fecal indicator bacteria (FIB) assays. We selected a subset of the mesocosm samples (two environmental and two enteric strains; Table 1) for total community RNA sequencing with an internal spiked control for absolute transcript quantification. Because the original RNA extraction protocol used was designed for rRNA:rDNA analysis and was not optimized for mRNA sequencing (i.e., depletion of 16S rRNA gene transcripts), we were not able to get enough total RNA for rRNA-subtracted libraries. Thus, total RNA was sequenced instead. The resulting metatranscriptome libraries yielded an average of 3.2×10^7^ (± 9.7×10^6^) reads per sample and ~95.7% of those reads were rRNA. The internal RNA standard recovery in each metatranscriptome ranged from 0.02% to 0.13% of the original spike-in quantity, as originally planned. The internal standard percent recovery was used to estimate the absolute number of mRNA reads per ng of RNA sequenced (5.9×10^7^ ± 3.5×10^7^ on average in each sample) following the methods described in Satinsky et al. 2013 (23).

Reference genome sequences of the isolates were previously determined (Table 1) and were used for read mapping to identify genes with significantly different expression between the two isolate types. Overall, there were no differentially expressed genes (DEGs) between enteric and environmental isolates across all time points. When controlling for the effect of time, there were only 31 strain-specific DEGs with P_adj_ < 0.05 observed between D1 and D3, with 24 and 8 genes being more expressed in the environmental and enteric isolates, respectively (Figure 2). None of these genes were among the habitat-specific genes identified by the previous comparative genomic studies, such as the nickel uptake operon, *nik(MN)QO* (4–6). The DEGs found in the environmental isolates were mostly housekeeping genes such as ribosomal and transcription-related proteins (e.g., tRNA ligase and elongation factor T; Figure 2). In contrast, genes potentially related to cellular stress response, such as a putative transcription repressor (*niaR*), a DNA replication and repair gene (*recF*), and a zinc-transporting ATPase (*zosA*) had higher expression in the enteric isolates. Metal ions, such as Zn^+2^, Cu^+2^, and Mn^+2^, are known to be important for oxidative stress defense in commensal *E. faecalis* (24). Together these results suggest that a few genes expressed differently among the strains may be linked to different habitat adaptation but the great majority of genes in the genome did not show differential gene expression.

**Figure 2:**
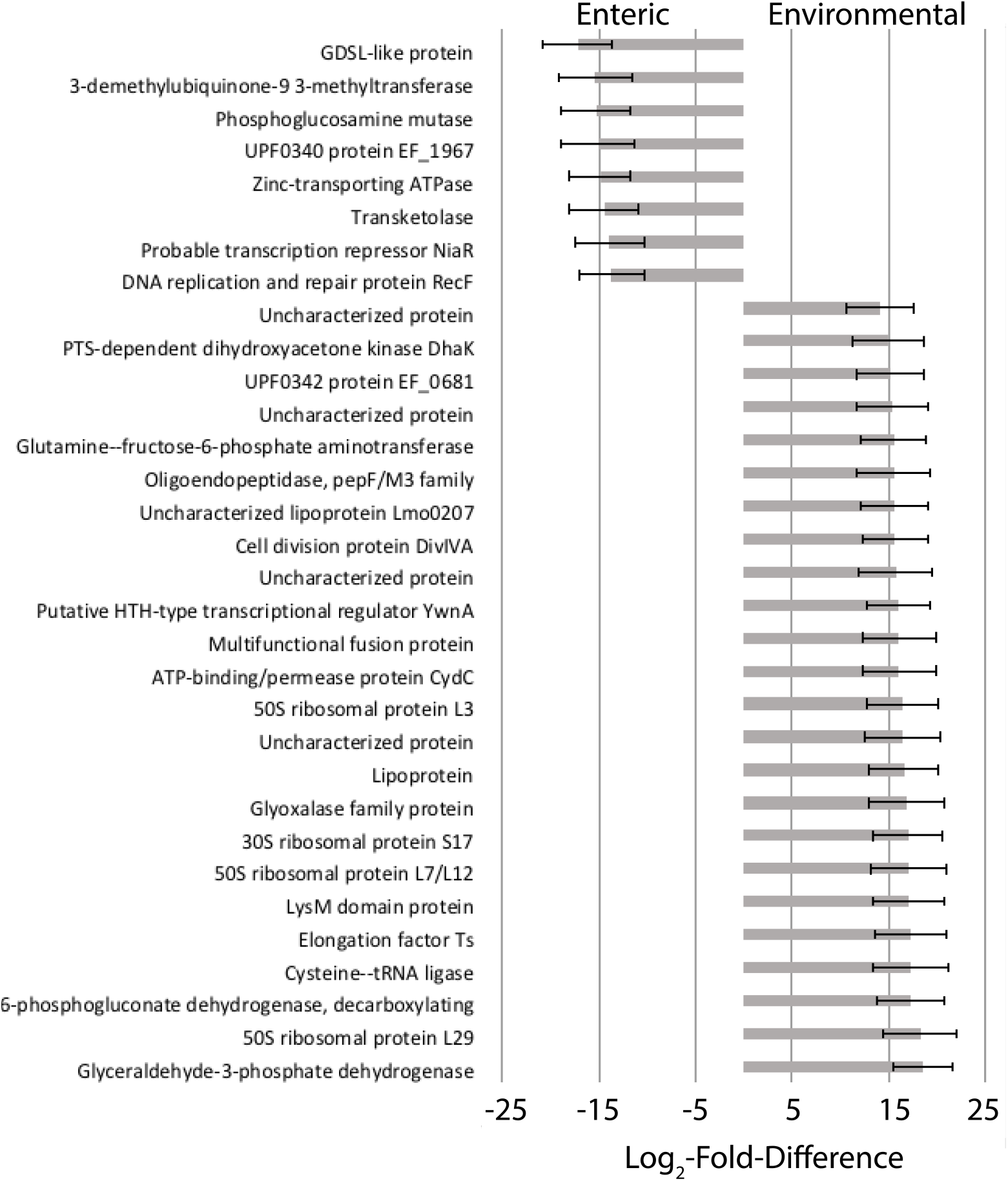
Differentially expressed genes between enteric and environmental *E. faecalis* isolates between days 1 and 3. Functional genes that were significantly more expressed (P_adj_ <0.05) in enteric relative to the environmental isolates (negative log_2_-fold-difference) or significantly more expressed in environmental relative to enteric isolates (positive log_2_-fold-difference). Error bars represent the standard error among biological replicates.

### rRNA:rDNA ratios over the standard growth curve in pure culture

Since the relationship between rRNA:rDNA ratios and growth rate are taxa-specific (16), we also collected baseline data on rRNA:rDNA levels in pure cultures of *E. faecalis* under standard laboratory conditions, which has not been examined previously for this species. A typical bacterium growth curve was observed in these experiments, in which the exponential growth phase lasted ~10 hours and maximum cell density (1.5×10^9^ CFU/mL) was observed at 12.5 hours (Figure 3). Cell density remained relatively stable until the next measurement at 25 hours, at which time cell density was still around 1.1×10^9^ CFU/mL (Figure 3A). The rRNA:rDNA ratios ranged from 5.5 to 372, with the lowest ratios being observed during early exponential growth phase (i.e., during the first 5 hours), after which point the ratio started to increase but there was a high level of variation between biological replicates (Figure 3B). The highest levels of rRNA:rDNA ratios were observed in the early stationary phase (~hour 12; average ratio = 372), after which the ratios started to decrease during stationary and death phases.

**Figure 3:**
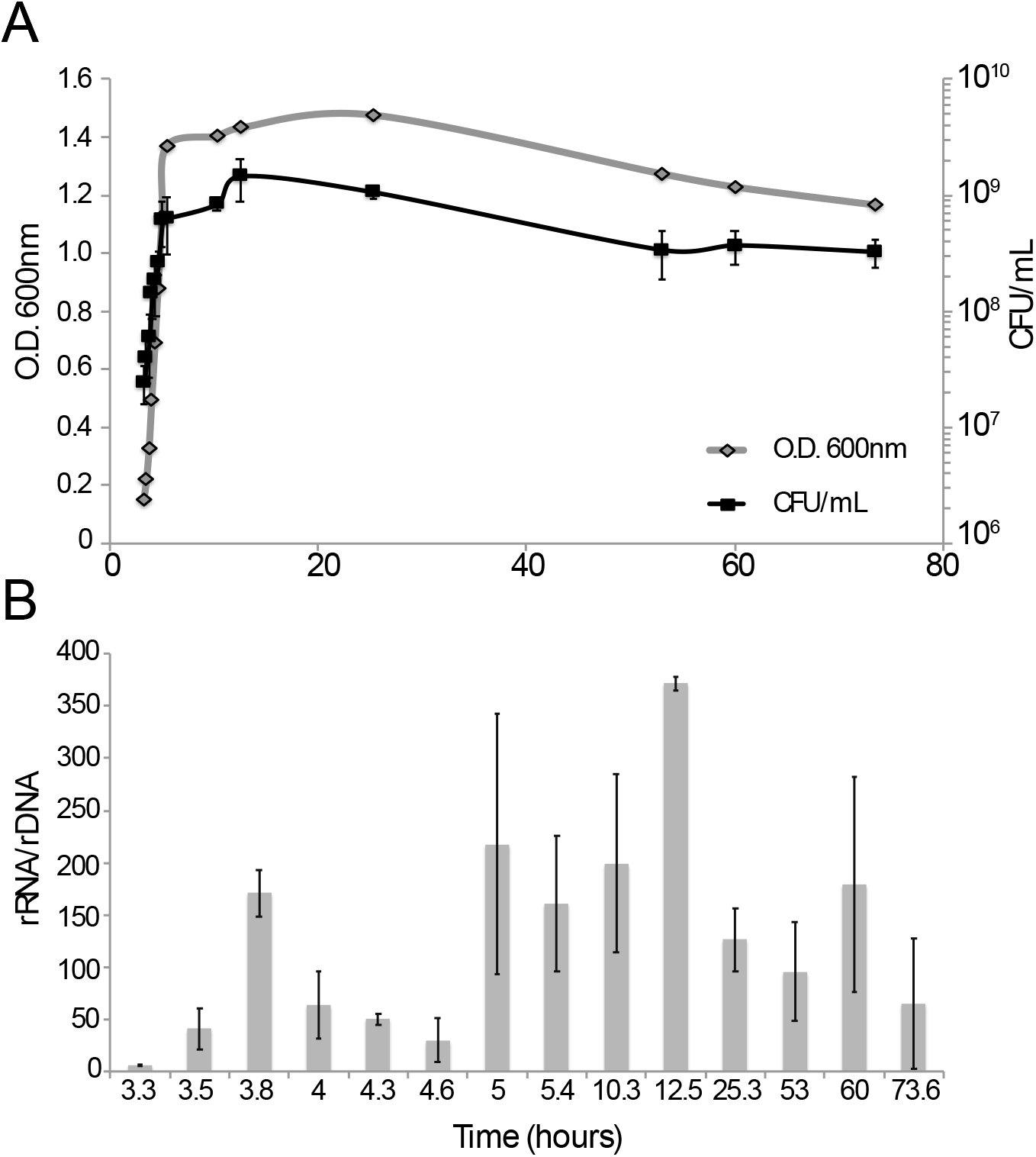
**(A)** Cellular abundance and **(B)** rRNA:rDNA ratios for *E. faecalis* MTUP9 in triplicate batch pure culture conditions. Error bars are standard deviation of biological and technical replicates.

## DISCUSSION

This study investigated whether the rRNA:rDNA ratio can be used to distinguish enteric versus environmental strains of *E. faecalis* for improved environmental water quality monitoring (20). We observed high variability in rRNA:rDNA ratios among biological replicates for all strains under oligotrophic mesocosm growth conditions (Figure 1B). Notably, the ratios under these conditions were, on average, roughly two orders of magnitude lower than those observed under standard lab conditions in pure culture (Figure 3B). These results are consistent with another study, which showed a high standard deviation in ratios and that copiotrophs have much lower ratios during growth in oligotrophic conditions relative to growth in rich media (25). Therefore, it appears that the rRNA:rDNA ratio could reflect oligo- vs. copio- trophic growth conditions for *E. faecalis*.

Although there was a difference between rRNA:rDNA ratios observed in the enteric and environmental isolates on D3 (Figure 1B), our results were not conclusive with respect to whether or not this assay is suitable for distinguishing isolate types in water quality monitoring applications because the differences were not large enough and were strain-specific (as opposed to habitat-type-specific), at least for the conditions tested here. However, the rRNA:rDNA ratio may be useful in pinpointing the age of a fecal pollution incident. All six isolates had significantly higher rRNA:rDNA ratios on D1 and D3 compared to D8 and D11, with overall average ratios of 11.7, 24.8, 4.7 and 1.8, respectively (Figure 1B). That is, the rRNA:rDNA ratio was substantially higher in the early stages, and this could serve as a sign of recent fecal pollution. Specifically, higher ratios (~12; based on the median rRNA:rDNA ratios observed for all strains on D1 and D3) could indicate a more recent pollution event, whereas lower ratios (e.g., <1.5; based on the median ratios observed for all strains on D8 and D11) could indicate that the public health risks from exposure to pathogens are not as high. The rRNA:rDNA can also reflect the physiological status of the group, as a whole, in environmental waters (20) and thus, may also be explored further as a tool for determining favorable conditions for regrowth from the VBNC state.

The viable cell counts indicated that the abundance of *E. faecalis* still exceeded the EPA recreational water quality criteria of 36 CFU/100 mL for all isolates on D8 (~10^5^ CFU/mL; Figure 1A); thus, these lake water samples would still be considered a public health risk according to current EPA standards (26). However, our findings that the rRNA:rDNA ratio decreases after D4 suggests that these cells have largely become inactive (e.g., enter VBNC) and/or have started dying; hence they represent a lower risk compared to cells at D1. Consistent with these interpretations, a recent quantitative microbial risk assessment (QMRA) analysis of sewage pollution suggested that the risk of exposure to pathogens is not significant after three days (27). In water bodies that consistently exceed EPA regulations for enterococci, it could be useful to investigate whether this is the result of a natural reservoir (i.e., no pathogen risk) or chronic pollution (pathogen risk) and techniques like the rRNA:rDNA assay presented here could be useful to help inform appropriate monitoring, management and/or mitigation strategies.

Notably, two of the three enteric isolates showed a six-fold increase in their rRNA:rDNA ratios from D1 to D3 (Figure 1B) and the ratios on D3 for these two isolates (~45 rRNA:rDNA) were similar to the lower end of average values observed for *E. faecalis* in pure culture (e.g., during early exponential phase; Figure 3B). However, our results based on viable cell or PCR counts suggested that *E. faecalis* isolates were not actively growing or replicating in the mesocosms over time. Moreover, the incubation conditions are remarkably different between oligotrophic growth and growth in pure culture, hence the trends observed in the two experiments (i.e., lake water mesocosm vs. pure batch culture) are presumably the result of different biological factors.

A potential explanation for increasing rRNA:rDNA ratios coupled to the decreasing cell counts observed in the oligotrophic mesocosm conditions is that the enteric isolates are increasing gene expression for pathways related to non-growth activities, such as environmental stress or cell homeostasis that results in more ribosomes (and thus more rRNA). Accumulating or maintaining high rRNA levels during periods of low activity may confer a competitive advantage upon return to favorable conditions, especially in copiotrophic environments that favor fast growers that can respond quickly to nutrient stimuli (28). Enteric isolates maintaining high cellular rRNA levels through D3 could indicate an adaptive strategy for high nutrient environments like the gut, whereas the environmental isolates are not “evolutionarily primed” to expect high nutrient influxes and do not devote as much energy to maintain high rRNA levels.

Consistent with these interpretations, metatranscriptomics analysis of the mesocosm incubation samples revealed that the enteric isolates differentially expressed several genes (DEGs) whose functions potentially reflect a stronger stress response compared to environmental isolates. This conclusion is also supported by a recent study that showed soil microbes adapted to low phosphorus conditions had much higher transcription of housekeeping genes under phosphorus-limitation (29). In contrast, DEGs of environmental strains included only a few housekeeping genes such as ribosomal and transcription-related proteins (Figure 2), which may indicate better survival because they are able to maintain general gene expression without a strong signal of environmental stress. However, the number of DEGs overall was small (only 31 DEGs in total and 30 of these had P_adj_ > 0.031) and these results may be spurious as about half of these DEGs detected could be due to chance based on the false discovery rate predicted by the DESeq2 analysis (i.e., expected ~17 DEGs by chance). Therefore, although our results provided some evidence that environmental and enteric isolates may respond differently upon release to the natural environment, the differences observed were too small and/or not consistent enough to provide robust means to distinguish between these two groups of isolates, at least for the conditions simulated by our mesocosm. It is possible that the environmental isolates tested here are not truly adapted to grow in the extra-enteric environment or -at least- the specific conditions of our mesocosms (e.g., they quickly died out during our mesocosm) and this accounted, partly, for the overall small differences observed with enteric isolates. However, the environmental isolates were obtained during regular monitoring of watersheds using the established EPA methods (4); thus, our results are relevant for microbial water quality applications, in any case.

Previous studies of other copiotrophs under balanced growth conditions in pure culture have shown that cellular rRNA concentration correlates well with growth rate (13, 30, 31). As such, we expected to see the highest rRNA:rDNA ratios for *E. faecalis* during the exponential phase in pure culture. However, the highest ratios were observed around hour 12 when growth was entering early stationary phase (Figure 3). The relationship between RNA levels and growth is not linear or consistent between different taxa, especially in environmental oligotrophic bacteria (18, 32). Therefore, this result is not necessarily surprising, but does suggest that the regulation of cellular RNA levels in environmental isolates of *E. faecalis* may be more complicated and not linearly correlated to growth. One possible explanation for the trends observed is that during early exponential phase, the cells are rapidly replicating their genomes and may have multiple genome copies per cell during rolling replication, resulting in the observed lower rRNA:rDNA ratios. As nutrients in the batch culture start to become depleted and cell growth slows, there is a lag in the ribosome transcription feedback loop around hour 12 at which time ribosome concentration briefly exceeds cell demand for rapid growth and results in the observed higher ratio. The high variation between biological replicates observed in both experiments also suggests that this rRNA:rDNA assay should be tested in more isolates in order to confirm the preliminary trends reported herein and the amount of natural variability in rRNA:rDNA ratios between isolates as well as to provide more support for the explanations proposed above.

Furthermore, we acknowledge that the growth conditions employed in this study may also limit our ability to distinguish isolates from the two habitat types. Previous starvation experiments showed that in some taxa, growing cells at maximum or medium growth rates before starvation can affect whether high rRNA levels are sustained even when cell activity decreases (33, 34) and suggested that an organism’s response to an event (e.g., introduction to extra-enteric environment through fecal shedding) can be determined by the existing conditions before that event. In our dialysis bag mesocosm experiment, we spiked pure cultures grown in rich media into lake water, which may not accurately reflect the life histories of environmental or enteric *E. faecalis* isolates, and thus different ratios may be observed *in situ* relative to our mesocosm condition. For example, an enteric cell is likely first introduced into a sewage or septic system which may not be nutrient limiting but have other stressors like oxidation or predation before reaching a surface water body. It is also possible that the habitat-specific genes previously identified such as those encoding the nickel and cobalt transport systems in the environmental genomes (4–6) are tuned for different conditions or stimuli rather than the mesocosm conditions used here and this accounts for the lack of differential expression of these genes in our datasets. Although mesocosm studies are helpful for comparing *E. faecalis* survival in a more controlled environment, they cannot simulate all of the complex biotic and abiotic factors that occur in aquatic habitats. Inspecting the ratios in extractions directly from known, natural extraenteric reservoirs of *Enterococcus* such as in algal mats (35) could help to get a better understanding of how rRNA levels are regulated in isolates that have been (presumably) under nutrient limitation for a longer period of time. For example, *E. faecalis* rRNA:rDNA ratios observed in water samples after a combined sewer overflow (CSO) event were ~1.85 (20), which is similar to the ratios observed on D11 here, but it is not clear how long the CSO *E. Faecalis* populations were exposed to the extra-enteric environment and how this rRNA:rDNA ratio relates to public health risks.

Finally, the RNA extraction protocol used in this experiment was originally optimized for the rRNA:rDNA assay (i.e., simultaneous and consistent extraction of both DNA and RNA from a single filter to ensure that the same amount of starting material is used for both qPCR and RT-qPCR) and resulted in extractions with total RNA concentrations too low for rRNA-subtracted libraries. Accordingly, our metatranscriptomics datasets included a minority of reads representing protein-coding genes (< 5% mRNA; typical for non-rRNA-subtracted libraries) and some of the DEG signal could have been lost as a result of this (i.e., only the most highly expressed transcripts were detected in the metatranscriptomes due to low sequencing coverage of mRNAs overall). It should be mentioned, however, that cDNA reads covered the whole *E. faecalis* references genomes at 9x, on average for D1, thus, we should have been able to detect most DEGs on D1. In later time points, when the *E. faecalis* metatranscriptome signal was decreasing, consistent with the decreasing viable cell counts, we were able to detect only highly expressed genes as DEF based on a ~2X coverage of the genome, on average, by cDNA reads. Thus, even though a few truly DEG may have escaped detection for the reasons mentioned above, the overall small differences observed in the transcriptomes of enteric vs. environmental isolates represent a robust result. Future studies should include separate RNA extractions for the rRNA:rDNA ratio assay and metatranscriptomic sequencing.

Despite these limitations, our work provides useful information on rRNA:rDNA ratios in *E. faecalis* under both standard lab and *in situ*-like conditions relative for environmental water quality monitoring, which has only been investigated in natural water samples for this genus (20, 36). Our results provide some evidence for different habitat adaptations between environmental and enteric strains but the difference may be too subtle or not consistent enough to be used in water quality monitoring. Further, working with RNA is generally more difficult and expensive compared to DNA (e.g., often requires a −80 °C freezer, RNase-free consumables, etc.) and requires more technical expertise and higher sterility making this approach impractical for local municipalities or regulatory agencies with limited laboratory resources. That said, our study showed that the rRNA/rDNA ratio may be useful for determining more recent fecal vs. older pollution events. Furthermore, our results provided new insights on the relationship between rRNA levels and non-growth activities, such as in VBNC cells, for an important FIB taxon. Evaluating more strains and growth conditions would be necessary to confirm our preliminary findings and establish whether rRNA:rDNA-based methods can provide more robust public health risk assessments.

## MATERIALS AND METHODS

### Comparison of rRNA:rDNA ratios of enteric vs. environmental *E. faecalis* isolates in dialysis bag mesocosms

#### Dialysis bag mesocosm set-up

Lake water was collected from Lake Lanier (Georgia, USA; 34° 15’ 38.898”N, 83° 56’ 56.0328”W) in June 2017 using acid-washed 10 L carboys and transported immediately back to the lab and stored at 4°C for mesocosm set-up the following day. Lake water used for inoculation with the *E. faecalis* isolates was first filtered through 0.2 μm sterivex filters as described previously (37) to remove predation pressure. The remaining unfiltered water was used to fill 10-gallon aquarium tanks in which dialysis bags were suspended during the incubations, as described below. Frozen glycerol stocks of the *E. faecalis* isolates (Table 1) were streaked for single colonies onto tryptic soy agar (TSA) plates and grown overnight at 37 °C. A single colony from each isolate was then inoculated into four mL of tryptic soy broth (TSB) and incubated at 37 °C with shaking at 150 rpm for 14 hours. One mL from each overnight culture was washed once with phosphate buffered saline (PBS) before inoculation into filtered lake water to a final concentration of ~10^6^ CFU/mL. The initial concentration for each overnight culture was also determined by plate counts on TSA plates. The dialysis bags (6-8 kDa molecular weight cutoff) were filled to a total volume of 110 mL (~21 cm length of 32mm diameter dialysis tubing) and closed on both ends using polypropylene Spectra/Por clamps (Spectrum Laboratories). The dialysis bags have a pore size that allows passage of small molecules and ions but prevents passage of bacterial cells and viral particles. Enough dialysis bags were filled to sample each isolate in triplicate at four time points, plus four filtered lake water negative control bags. The dialysis bags were then transferred to 10-gallon aquarium tanks filled with unfiltered lake water and stored in environmentally controlled rooms at 22 °C in the dark. A small water pump was included in each tank for aeration and nutrient distribution. A small headspace of air was left in each bag when sealing with the clamps so that they could float freely in the tanks.

#### Mesocosm sampling

Each sampling time point included triplicate biological replicates per isolate and a single lake water negative control. Destructive sampling of the dialysis bags occurred at days 1, 3, 8 and 11 after the initial set up. Fifty mL from each dialysis bag was filtered through 0.45 μm polycarbonate membranes, then the filters were transferred to 2 mL screw-cap tubes that had been pre-filled with 0.8 mL Qiagen buffer RLT (with 1% beta-mercaptoethanol) and 100 mg of acid-washed 0.1 mm glass beads. Bead tubes were stored at −80 °C until ready for extraction (storage time was <1 month for all filters). Additionally, water from each bag was 10-fold serially diluted with PBS for culture-based enumeration on TSA and mEnterococcus Agar (BD Difco™) plates. All dilutions yielding measurements within the acceptable range of quantification were averaged to estimate CFU/mL of each isolate. The filtered lake water was also checked for sterility by plating on TSA and mEnterococcus at day 0 before inoculating with *E. faecalis* isolates.

#### Total nucleic acid extraction

The frozen filters were defrosted on ice before the cells were mechanically lysed using a BioSpec Mini-BeadBeater-24 in four 1-minute intervals with icing in between to prevent the samples from excessive heating and to protect the integrity of the RNA. Total nucleic acids were extracted from cell lysates using the Qiagen AllPrep DNA/RNA mini extraction kit following the manufacturer’s protocol for animal tissue. Contaminating DNA was removed from RNA samples by digestion (1-2 times depending on the sample concentration) with the Ambion TURBO DNase kit following the manufacturer’s protocol. RNA integrity was assessed with an Agilent 2100 Bioanalyzer instrument and the Agilent RNA 6000 Pico kit. RNA samples used for downstream analysis generally had a RIN >7 and 23S/16S rRNA ratio >1.

#### Assessment of the quality of the RNA and DNA extractions

Elimination of DNA from RNA samples was confirmed by end-point PCR amplification with the same primers used for the *E. faecalis* specific 16S rRNA qPCR assay (38). Two μL of undiluted RNA was used as template in 20 μL PCR reactions with 0.5 μM primers, 200 μM dNTPs, 0.5 units TaKaRa Ex-TAQ polymerase and 1x TaKaRa PCR buffer. The thermocycling conditions are as follows: 1 min at 95 °C then 30 cycles of 95 °C for 15 sec and 61 °C for 30 sec followed by 72 °C for 1 min. The PCR products were visualized with gel electrophoresis and the absence of any detectable bands in the gel indicated that there was no significant DNA contamination in the RNA samples.

The absence of PCR inhibitors in the RNA samples was confirmed by end-point PCR amplification in which a known amount (~10^7^ copies) of standard plasmid DNA was spiked into a PCR reaction in the presence of RNA. The same PCR master mix and thermocycling conditions were used as above except the primers targeted the nickel uptake gene in the standard plasmid (not published). If the RNA preparation contained inhibitors, amplification of the DNA template was expected to be inhibited in the presence of RNA. The PCR products were run on a 1% agarose gel and the presence of a single band at the expected size and yield of the PCR amplicon from the standard plasmid template confirmed the absence of any PCR inhibitors.

#### Quantification of 16S rRNA and rDNA using reverse transcriptase quantitative PCR (RT-qPCR) and quantitative PCR (qPCR)

DNA and RNA concentrations were quantified using the Qubit High Sensitivity DNA and RNA kits (Thermo Fisher Scientific), respectively, and the Qubit 2.0 fluorometer. Template nucleic acids were then diluted to below 0.5 ng/μL before amplification using an *E. faecalis* specific 16S rRNA gene assay (38). The standard plasmid used for absolute quantification was a full-length *E. faecalis* 16S rRNA gene ligated into the pCR™2.1-TOPO^®^ TA vector and cloned into One Shot^®^ Chemically Competent TOP10 *Escherichia coli* using the TOPO^®^-TA cloning kit (Invitrogen) following manufacturer’s instructions. Eight 10-fold serial dilutions (10^8^ to 10^1^ copies per reaction) of qPCR standard plasmids were assayed in triplicate on every 96-well plate for absolute quantification. All reactions were performed on the Applied Biosystems 7500Fast machine using Bio-Rad iTaq™ Universal Probes One-Step reagents following the manufacturer’s protocol. Reactions were performed in triplicate in a total volume of 20 μL that included 2 μL of the template or standard plasmid and 250 nM of each primer and TaqMan (5’ hydrolysis) probe (the RT-qPCR reactions also included 0.5 μL of Bio-Rad iScript advanced reverse transcriptase). Thermocycling conditions for qPCR consisted of an initial 50 °C step for 2 min followed by 95 °C for 10 min, then 40 cycles of 95 °C for 15 sec and 60 °C for 60 sec. The RT-qPCR thermocycling conditions were the same except for the initial step of 50 °C for 10 min followed by 95 °C for 2 min. The calibration curve from each plate was used to calculate rRNA and rDNA copy numbers in each sample which were averaged among technical replicates, multiplied by elution volume (200 or 50 μL for DNA and RNA, respectively), and then divided by the filter volume (50 mL) to give total copies per milliliter of mesocosm water sampled.

#### Statistical analyses

Culture-based cell counts over time were averaged by habitat type and tested for significant difference between the two groups using the paired Wilcoxon signed rank test in base R. The rRNA:rDNA ratios of each isolate were compared by habitat type overall and between each time point. Since the data violated the Bartlett test of equal variance (as implemented in base R), nonparametric pairwise multiple comparisons were performed using the Wilcoxon signed rank test with the Holm p-value correction for multiple comparisons using a custom R function (http://www.statmethods.net/RiA/wmc.txt).

### Metatranscriptome sequencing and analysis of total RNA from dialysis bag mesocosms

#### Metatranscriptome library preparation and sequencing

Triplicate RNA extractions from each isolate at each of the four time points were pooled together in order to obtain enough high-quality RNA for metatranscriptomic sequencing, and cDNA libraries were prepared using the ScriptSeq v2 RNA-Seq Library Preparation kit (Illumina) following the manufacturer’s instructions except a half ng (~1% of total library size) of a luciferase internal RNA standard was included during the RNA fragmentation (step 3A) for absolute quantification of transcript copy numbers as described below. The quality and insert size of each cDNA library was determined using the Agilent High Sensitivity DNA kit and Agilent 2100 Bioanalyzer instrument. Library concentrations were determined using the Qubit HS DNA kit before pooling and loading into the flow cell according to manufacturer’s recommendations and sequencing on the Illumina HiSeq 2500 instrument as described previously (29).

#### Luciferase internal RNA standard preparation

The Promega pGEM^®^-luc plasmid vector (accession number X65316) containing a 1094 nucleotide fragment of the firefly luciferase gene was digested with SphI-HF restriction enzyme (New England Biosystems) at 37 °C for 1 hour to linearize the plasmid, followed by the Qiagen PCR clean-up kit to stop the reaction. The digested DNA was gel-purified using 1.5% low melt agarose gel and the MO BIO UltraClean^®^ 15 DNA purification kit followed by end repair with the Thermo Scientific Fast DNA End Repair kit and another clean up with the Qiagen PCR clean-up kit but with a 30 μL elution volume. The DNA was concentrated by ethanol precipitation before transcribing to RNA with the Promega Riboprobe^®^ *in vitro* Transcription T7 System and following the manufacturer’s protocol 4.F for synthesis of large amounts of RNA. The RNA standard quantity and quality were determined using the Qubit HS RNA kit and Agilent Bioanalyzer as described above.

#### Transcriptome sequence analysis

All transcriptomic reads were quality filtered and trimmed as described previously (39). Trimmed reads were filtered to remove rRNA sequences using SortMeRNA v2.1 (40) with all rRNA databases in the program and the following options: --blast 1 --num_alignments 1 -v -m 8336. The internal luciferase standard sequences were identified by blastn search against the 1094 bp length nucleotide luciferase reference sequence carried on the pGEM^®^-luc plasmid vector (accession number X65316). Luciferase matches were filtered for best match using a threshold of 97% identity and alignment length that is 80% of the query read length, and resulting matches were subsequently removed from the transcriptomic datasets. The number of internal standard sequences recovered was used to estimate the absolute number of mRNA transcripts in the sample and sample sequencing depth was defined as the actual number of mRNA reads sequenced in metatranscriptome divided by the absolute number of mRNA transcripts in the sample (23). Metatranscriptomic short reads have been deposited to the NCBI SRA database under BioProjectID PRJNA720051.

Reference genome assemblies for the four isolates that were used as inocula in the mesocosms were downloaded from NCBI (Table 1). Prodigal v2.6.1 (41) was used to predict genes from the assemblies, which were then annotated against the Swiss-Prot database (downloaded March 2019; (42)) using blastp (options: evalue 1E-6 and max_target_seqs 10; (43)). Matches to the reference Swiss-Prot sequences were filtered for best matching, using 40% identity and 40% query cover alignment length as threshold values. All genes that had no match to the Swiss-Prot database were annotated against the TrEMBL database (downloaded May 2018; (42)) using the same match filtering cut-off. Non-rRNA metatranscriptomic reads (i.e., after removing internal standard sequences) were mapped against predicted genes using MegaBLAST (43) for the corresponding isolate that was used as an inoculum in that sample and matches with <97% identity and <50 bp alignment length were removed from further analysis. Read count tables against predicted genes were generated using custom scripts and were used as the input for DESeq2 v1.16.1 (44) and the sample sequencing depth as determined from the internal standard was used for the estimate size factors step. Differentially expressed genes between enteric and environmental isolates were determined using the Likelihood Ratio Test and false discovery rate (P_adj_ <0.05) as implemented in DESeq2.

### rRNA:rDNA ratio of *E. faecalis* in pure culture under standard laboratory conditions

#### Batch culture growth conditions and sampling

*E. faecalis* strain MTUP9 (Table 1) was streaked for single colonies onto a TSA plate and grown overnight at 37 °C. A single colony was then inoculated into 4 mL of TSB and incubated at 37 °C with shaking at 150 rpm for 14 hours. The overnight liquid culture (100 μL) was inoculated into 60 mL fresh TSB in triplicate cultures to start the growth curve experiment (O.D._600_ < 0.1 at time 0) and incubated at 37 °C with shaking at 150 rpm. Each triplicate culture was sampled at 14 time points over 73 hours to capture the different growth phases. At each sampling point, 1 mL of each triplicate culture was collected for O.D._600_ reading, 0.1 mL was serially diluted 10-fold in PBS for plate counts on TSA, and 0.5-1 mL of the culture was collected for nucleic acid extraction by centrifuging at 10,000 rpm for 5 minutes and decanting the supernatant. Cell pellets were re-suspended in 600 μL buffer RLT (Qiagen) with 1% beta-mercaptoethanol and stored at −80 °C until ready for extraction. The re-suspended cell pellets were defrosted on ice and transferred to 2 mL screw cap tubes pre-filled with 100 mg of acid-washed 0.1 mm beads. Total nucleic acids were extracted and used for rRNA:rDNA analysis following the same protocol for filters as described above.

## ACKNOWLEDGMENTS

This work was supported by the US National Science Foundation, award numbers 1511825 (to J.B. and K.T.K.) and 1831582 (K.T.K.), and the US National Science Foundation Graduate Research Fellowship under grant number DGE-1650044 (to B.S.). These funding agencies had no role in the study design, data collection and analysis, decision to publish, or preparation of the manuscript. The research presented was not performed or funded by EPA and was not subject to EPA’s quality system requirements. The views expressed in this article are those of the author(s) and do not necessarily represent the views or the policies of the U.S. Environmental Protection Agency.

